# Interactively Integrating Reach and Grasp Information in Macaque Premotor Cortex

**DOI:** 10.1101/2024.06.12.598592

**Authors:** Junjun Chen, Guanghao Sun, Yiwei Zhang, Weidong Chen, Xiaoxiang Zheng, Shaomin Zhang, Yaoyao Hao

## Abstract

Successful reach-to-grasp movements necessitate the integration of both object location and grip type information. However, how these two types of information are encoded in a single brain region and to what extend they interact with each other, remain largely unknown. We designed a novel experimental paradigm that sequentially prompted reach and grasp cues to monkeys and recorded neural activity in the dorsal premotor cortex (PMd) to investigate how the encoding structures change and interact during arm reaching and hand grasping movements. This paradigm required monkeys to retain the first prompted cue when the second one arrived, and integrate both to accomplish a final goal movement. PMd neurons represented both reach and grasp to similar extend, yet the encodings were not independent. Upon the arrival of second cue, PMd continued to encode the first cue, albeit with a significantly altered structure, as evidenced by more than half of the neurons displaying incongruent modulation. At a population level, the encoding structure formed a distinct subspace that differed from, but was not entirely orthogonal to, the original one. Employing canonical correlation analysis, we identified a subspace that consistently preserved the encoding of the initial cue, potentially serving as a mechanism for downstream brain regions to extract coherent information. Furthermore, this shared subspace comprised a diverse population of neurons, including both congruent and incongruent units. these findings support the argument that reach and grasp information are interactively integrated within PMd, with a shared subspace likely underpinning a consistent encoding framework.

## Introduction

Primates, including humans and monkeys, frequently engage in reach-to-grasp movements to manipulate objects within their environments. These movements entail the intricate coordination of reaching and grasping actions, which are respectively controlled by proximal and distal muscles (Castiello, 2005; Castiello and Dadda, 2019). Despite extensive anatomical and behavioral research on reach-to-grasp movements, their underlying neural coding mechanisms are notably complex and a subject of ongoing debate (Abbaspourazad et al., 2021; Alstermark and Isa, 2012; Suresh et al., 2020). On one hand, topological mapping between brain cortices and effectors suggests that reaching and grasping movement are controlled by distinct cortical networks (Jeannerod, 1984; Penfield and Boldrey, 1937; Whishaw and Karl, 2014). On the other hand, the high degree of coordination between these movements hints at deeper neural interconnections that extend beyond mere independent modulation (de Vries et al., 2018; Gordon et al., 2023; Graziano et al., 2002).

Historically, reaching and grasping movements have been considered as two separate categories and have been studied in isolation within neurophysiological research. Commonly, subjects are directed to reach to multiple positions with a uniform grip type (Churchland et al., 2012; Moran and Schwartz, 1999; Sergio et al., 2005) or grasp various objects at a fixed location (Hendrix et al., 2009; Saleh et al., 2010; Spinks et al., 2008).The dual parallel parieto-frontal circuits model has been the prevailing theory for the brain control of upper limb movements for the past decades (Luppino et al., 1999; Rizzolatti and Matelli, 2003; Tanné-Gariépy et al., 2002). According to this model, primate reaching movements are managed by a dorso-medial circuit, processing sensorimotor information from posterior parietal area V6A and medial intraparietal area (MIP) to dorsal premotor area (PMd). Meanwhile, the grasping movements are tuned in a ventro-lateral parieto-premotor circuit, involving anterior parietal areas AIP, PFG and ventral premotor area (PMv). Both pathways ultimately converge on the primary motor cortex (M1) to exert executive control. Although this classical model has garnered supports from various studies, including electrophysiology (Cui and Andersen, 2007; Raos et al., 2006), neural imaging (Cavina-Pratesi et al., 2010; Konen et al., 2013), tract-tracing (Gamberini et al., 2009; Gerbella et al., 2011) and lesion (Hwang et al., 2012; Vesia et al., 2010), some researches have presented divergent evidences challenging the independent coding theory for reaching and grasping, particularly concerning the dorso-medial circuit. For instance, the V6A area has been shown to activate when subjects are presented with objects to be grasped in functional magnetic resonance imaging (fMRI) studies (Maratos et al., 2007).

Furthermore, neurons responsive to reaching movements have been documented in the ventral pathway V6A (Breveglieri et al., 2016; Fattori et al., 2010) and neurons responding to grasping were recorded in dorsal pathway PMd (Raos et al., 2004; Stark et al., 2007). Additionally, lesions in the V6A area of monkeys have resulted in impairments not only to reaching movements but also to grasping and wrist orientation (Battaglini et al., 2002).

While PMd plays a pivotal role in reach-to-grasp movements, the extent of mutual influence between reach and grasp encoding is not well understood. Recent studies integrating multiple target positions and grip types have demonstrated that single neurons in premotor cortex could encode reaching and grasping movements concurrently (Hao et al., 2017; Rouse and Schieber, 2016; Takahashi et al., 2017). However, the detailed cooperative coding rules between these two movements remain to be elucidated. Firstly, it is unknown whether reaching and grasping movements are modulated independently or dependently in a hybrid coding neuron. For example, previous research has demonstrated context-dependent coding of hand grasping in monkey’s AIP neurons, where these neurons remained silent upon receiving grip type cues but became highly activated once grasp target was revealed (Baumann et al., 2009). Moreover, the cooperative coding involves not only the interaction when the two types of information are processed simultaneously in a cortex, but also the sequential dependence of one on another. Sequentially prompted sensory stimuli exhibit rich dynamics in the premotor cortex (Rossi-Pool et al., 2016; Rossi-Pool et al., 2017). Various types of information were integrated sequentially in prefrontal cortex (Rao et al., 1997; Tang et al., 2020), with modulations being temporally dependent (Cisek and Kalaska, 2005). Therefore, it is intriguing to explore whether the reaching position serves as a precondition to grip type coding and vice versa.

In this study, we recorded single-unit activity from PMd area while monkeys performed a sequentially prompted reach-to-grasp task with delays. The task involved reaching to one of two target objects with either a power or precision grip. To ascertain the sequential dependence of reaching and grasping modulation, we presented target position cue and grip type cue separately. The findings indicated that PMd neurons encoded both reach and grasp, yet the encodings were not independent. Upon the arrival of the second piece of information, PMd continued to encode the first but with a substantially altered encoding structure. A multivariate statistical approach captured the subspace that preserved consistent representation, potentially serving as the mechanism for congruent information extraction by downstream regions.

## Results

Two rhesus monkeys (*Macaca mulatta*, Monkey Y and W) were trained to perform a sequentially prompted reach-to-grasp (SPRG) task according to LED cues. In brief, the task involved reaching for and grasping one of two identical handles (left vs. right) fixed on a vertical panel in front of them (as depicted in Fig. 1A), utilizing two distinct grip types: power and precision. To investigate the interaction between reach and grasp encoding, the two types of information in SPRG were prompted to monkeys sequentially with delays, i.e., either reach position (‘pro’-mode) or grip type (‘anti’-mode) was showed up first and the other one followed after a fixed delay period (delay1). Post a subsequent random delay period (delay2), a go cue was issued and the monkey reached for the correct object and grasped it with correct grip type (Fig. 1B). This procedure required the subjects to retain the initial cue information during both delay1 and delay2, integrate the two cues during delay2, and formulate an accurate response.

**Figure 1.**
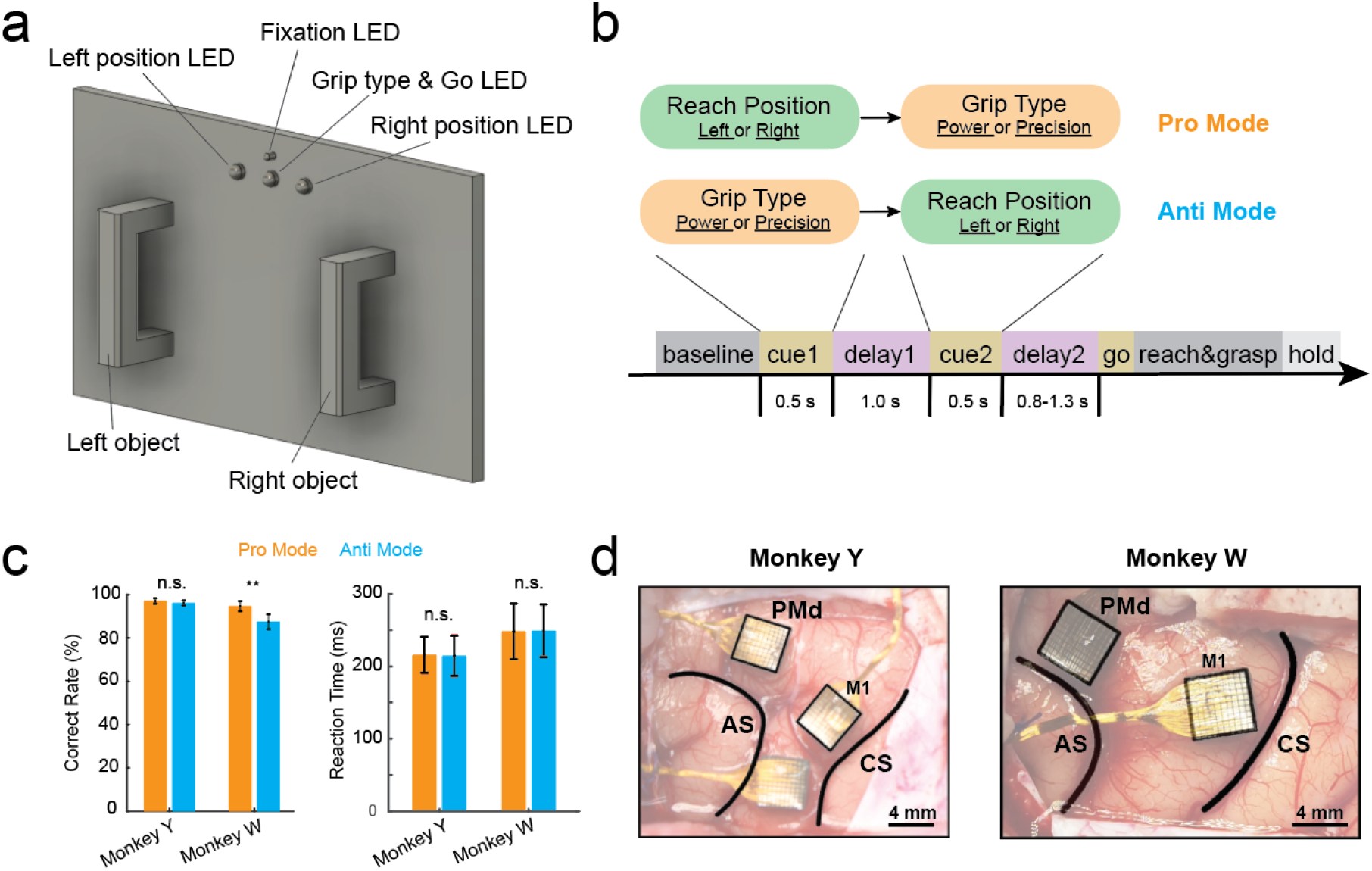
Behavioral task and electrode implantation. **(a)** Two identical handles are fixed on a panel in the left and right position for monkey grasping task. The panel is installed vertically in front of the subject. Two white LEDs are used to indicate which object to grasp (left position & right position LED). One central RGB-LED servers to prompt grip types (blue for power grip and yellow for precision grip) and Go cue (green). A small LED in the center is used for eye fixation. **(b)** Time line for the SPRG task. During Cue1, either reach position (pro-mode) or grip type (anti-mode) are prompted to the subject and the other information (i.e. grip type for pro-mode and reach position for anti-mode) was showed during Cue2 with 1-sec delay (delay1). After another delay (delay2), Go cue is issued and subject starts to grasp one of the objects with specific grip type and hold for reward. **(c)** Averaged behavioral performance (correct rate, left) and reaction time (right) for pro- and anti-mode trials for both monkeys. n.s., not significant; **, *p* < 0.01 (paired *t*-test). **(d)** The implantation sites of micro-electrode arrays for both monkeys. The thick black lines show the landmarks of Arcuate Sulcus (AS) and Central Sulcus (CS). Arrays in area of M1 and PMd are highlighted with black boxes.

Given the task’s design, there are four different combinations in each mode and a total of eight conditions with varied prompting sequences. The monkeys performed the SPRG task with pseudorandom trials of different modes and trial types within the same session, enabling the examination of the same neurons’ responses across both modes. The success rates of the monkeys, measured as percentages of correct reach-to-grasp movements, and reaction time, measured as the time between go cue and object contact, were highly consistent across two modes (with the exception of Monkey W, whose correct rates varied slightly between modes but remained above 85%) (as illustrated in Fig. 1C). This consistency suggests that the monkeys were equally adept at performing both modes on a trial-by-trial basis.

### PMd vs. M1 responses during SPRG task

We recorded single neuron activities from both dorsal premotor cortex (PMd) and primary motor cortex (M1) while monkeys carried out the SPRG task (Fig. 1D). M1 neurons were found less involved in the task than PMd neurons, responding only to the actual reach-to-grasp action without discrimination among different trial types (Fig. S1). First, the proportion of neurons encoding reach position and grip type in M1 was significantly lower than in PMd throughout the entire trial in both pro- and anti-mode, as was the representation of the interaction between reach and grasp (Fig. S1A). Secondly, the pattern of instantaneous population variance between M1 and PMd differs (Fig. S1B). The variance of M1 was almost flat during cue and delay epochs and peaked after go cue, whereas the variance of PMd fluctuated, peaking during cue2 when the integration of two pieces of information was required for the final motor plan. Lastly, we examined the neuronal connectivity between M1 and PMd (Fig. S1C), revealing that substantial connectivity occurred only after the go cue, when the subjects initiated the reach. Taken together, these data indicated that M1 activity observed here only responded to the gross reach-to-grasp movement, but not to any preparatory processes. Consequently, we only focused our subsequent analysis on the PMd, investigating how reach and grasp information are encoded and interact within this singular brain region.

### PMd encodes both reach and grasp

We recorded PMd activity across five sessions for each monkey as they performed the SPRG task. On average, 101 and 158 single units were isolated per session for Monkey Y and W, respectively. The vast majority of neurons exhibited significant modulation in response to either reach position or grip type during at least one epoch of the task. Given our interest in understanding how reach and grasp encoding interact and integrate within the PMd, we concentrated our analysis on the delay1 epoch, where only one piece of information was presented, the delay2 epoch, where the additional information was prompted and integrated, and the comparative analysis between these two epochs.

We recorded PMd activity across five sessions for each monkey as they performed the SPRG task. On average, 101 and 158 single units were isolated per session for Monkey Y and W, respectively. The vast majority of neurons exhibited significant modulation to either reach position or grip type during at least one epoch of the task. Given our interest in understanding how reach and grasp encoding are interacted and integrated in PMd, we concentrated our analysis on the delay1 epoch, where only one piece of information was presented, the delay2 epoch, where the additional information was prompted and integrated, and the comparative analysis between these two epochs.

To dissect the encoding properties of PMd for reach and grasp separately, we initially focused on the delay1 epoch, characterized by the provision of a single type of information—either reach position or grip type. Figure 2A presented the raster and peristimulus time histogram (PSTH) plots of two exemplar neurons during the task in both pro- and anti-mode. Neuron 1 demonstrates a clear distinction between left and right reach positions during the delay1 epoch in pro-mode, with less pronounced modulation for grip type in anti-mode. In contrast, Neuron 2 exhibits modulation in both pro- and anti-modes during the delay1 epoch, suggesting that this neuron encodes both reach and grasp simultaneously. Among the population, we quantified the proportion of neurons that significantly encoded position, grip or both. As depicted in Fig. 2B, the two monkeys revealed similar distributions across categories; an average of 21.5%, 17% and 38.5% of neurons significantly encode position, grip and both during delay1 epoch, respectively. We employed principal component analysis (PCA) to visualize this modulation in a reduced two-dimension space in Fig. 2C, which illustrated clear population level separation for position in pro-mode and grip in anti-mode. Furthermore, we sought to determine if the modulation strength for position and grip were comparable. Fig. 2D displays the distribution of modulation depth (MD, see methods) for all recorded neurons. The MD distributions were highly overlapped for position and grip encoding in both monkeys (quantification in Fig. S2D). This finding was corroborated by single-trial decoding analysis (see methods) shown in Fig. 2E, which shows that reach position in pro-mode and grip type in anti-mode could be accurately decoded from neural activity in delay1 epoch; however, the other information, i.e., grip type in pro-mode and reach position in anti-mode, were at the chance level. These data suggest that reach and grasp are equally encoded in PMd.

**Figure 2.**
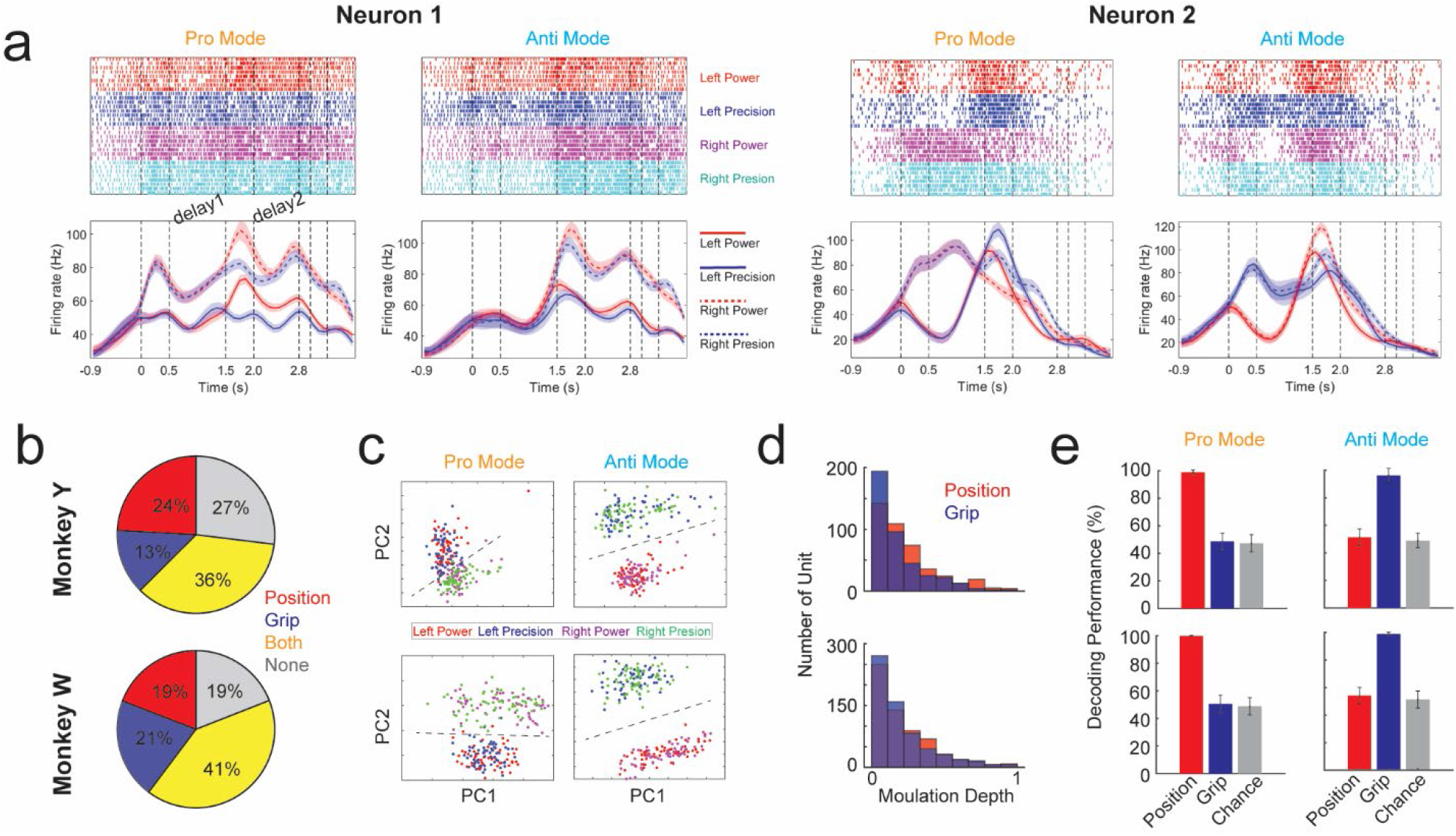
PMd neurons encode both reach and grasp movements during delay1. **(a)** The raster plot (upper) and averaged firing rate (lower) of two example neurons. The vertical dash lines separate the plot into 8 segmentations, which indicate baseline, cue1, delay1, cue2, delay2, move, reach & grasp, hold, respectively, as shown in Fig. 1B. Delay1 and delay2 are highlighted. The different colors in raster plot indicate different combinations of reach positions (left/right) and grip types (power/precision). In firing rate panel, solid (dash) lines indicate left (right) position; red (blue) color indicates power (precision) grip. **(b)** Percentage of neurons in PMd that modulate position (red), grip type (blue), both (orange) and neither (gray) during delay1. **(c)** Neural activity during delay1 was dimensional reduced to 2D space (PC1 vs. PC2) using principal component analysis (PCA). Each dot indicates averaged activity in one trial. Dash lines separate the reach position in pro-mode and grip type in anti-mode. **(d)** The histograms of modulation depth are plotted separately for position (red) and grip (blue), which shows similar distribution. **(e)** The decoding performances for position (red) and grip (blue) using delay1 activity in both pro- and anti-mode. The gray bar indicates chance level using shuffled data. Grip decoding in pro-mode and position decoding in anti-mode are not significantly different with the chance level (*p* > 0.05).

To further elucidate the reach and grasp encoding in PMd, we conducted an analogous analysis for delay2 epoch (Fig. S2). The percentage of neurons (Fig. S2A), PCA visualization (Fig. S2B) and decoding analysis (Fig. S2C) yielded results analogous to those of the delay1 epoch, indicating that reach, grasp, and their combination can be equally well distinguished. However, the MD was significantly higher for reach position in one of the monkeys during delay2 epoch. We then systematically compared the MD for reach and grasp in delay1, delay2 and move epoch (Fig. S2D), and only move epoch consistently showed significant higher reach MD for both monkeys. This suggests that PMd encodes reach and grasp with equal strength during memory and planning epochs but biases to reach when the actual movement is executed. This finding aligns with classical studies that position PMd as a key node in the dorsolateral reaching pathway.

### Reach and grasp are interactively integrated in PMd

To investigate if reach and grasp interact with each other in PMd, we compared neural activity in delay1 and delay2 to see if the arriving of the second information influences the encoding of the first information. First, we confirmed that PMd still encodes the first information as the second arrives during delay2, which is necessary for the subject to successfully complete the task (Fig. 3). The percentage of neurons that encode the first information remains consistent starting from the cue1 epoch until the end of move epoch (Fig. 3A). Quantification of the percentage in delay1 and delay2 epochs revealed a nearly equivalent number of neurons encoding the initial information (with a slight decrease observed in delay2 of the anti-mode for Monkey Y, 40.35% vs. 27.51%, *p* < 0.05) (Fig. 3B). Furthermore, we performed single-trial decoding analysis along the time course of trial to verify that the first information could be discerned during delay2 epoch. As shown in Fig. 3C, the first information was readily decodable from the cue1 epoch and remained so throughout the trial. Quantification in Fig.3D showed perfect decoding accuracy for both delay1 and delay2, with no significant difference between them. Concurrently, the four combinations of reach and grasp (chance level, 25%) were also perfectly decodable during delay2, contrasting with only around 50% during delay1.

**Figure 3.**
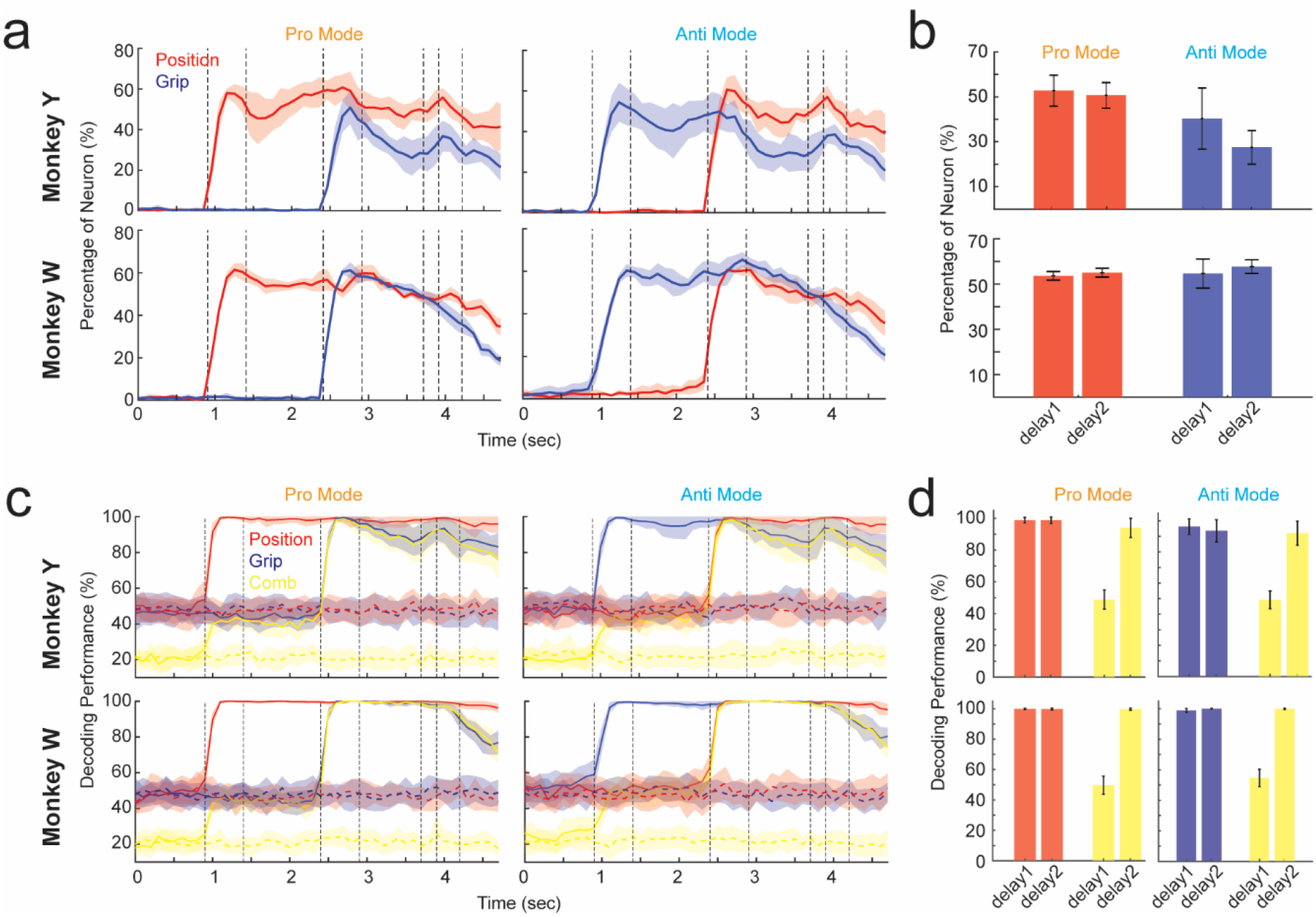
PMd still encodes the first information when the second one arrives. (a) Percentage of neurons that significantly modulate position (red) and grip (blue) are plotted against time during the task. The shadow area indicates standard deviation for different sessions. (b) Bar plot of percentage of neurons that modulate position (red) and grip (blue) during delay1 and delay2. Error bar indicates standard deviation. All comparisons are not significant (*p* > 0.05), except for anti-mode of Monkey Y (*p* = 0.0141). (c) Same as (a) but for decoding performance of position (red), grip (blue) and combination (yellow). The dash lines indicate chance levels calculated with shuffled data. (d) Same as (b) but for decoding performance during delay1 and delay2.

Next, we employed analysis of variance (ANOVA) to directly ascertain the proportion of variance in neural activity attributable to the interaction between reach and grasp, denoted as *eta* value. As outlined in the Method section, a non-zero *eta* value for interaction implies that reach and grasp do influence each other’s encoding. We presented the session averaged *eta* value for reach position, grip type and their interaction as a function of trial time in Fig. 4A. The patterns for position and grip coding were similar with the trend of percentage of neurons showed in Fig. 3A. Notably, the interaction emerged post cue2 onset, peaked around the offset of cue2, and faded away at the end of the trial. The quantification results (Fig. 4B) indicated a significant non-zero interaction component during delay2, alongside significant reach and grasp components. To elucidate various types of interaction between reach and grasp, three example neurons were depicted in Fig. 4C. Neuron 1 exhibits no interaction between reach and grasp (Two-way ANOVA test, *p* > 0.05). regardless of whether the monkey planned to reach left (solid) or right (dashed), Neuron 1 consistently prefers power grip (red) in delay2. The same holds true for anti-mode, where grip type does not alter the neuron’s preference for reach position in delay2. Neuron 2 and 3, however, displayed a significant interaction effect (Two-way ANOVA test, *p* < 0.001). For instance, in pro-mode, Neuron 2 favors precision grip (blue) over power grip (red) in delay2 when the monkey intends to reach the right position; should the reach position switch to left, the grip type preference vanishes. Neuron 3, in anti-mode, reversed the preference for reach position (solid vs. dash) based on the grip type (red vs. blue) prompted in delay1. The percentage of neurons with significant interactive effect was around 24% for Monkey Y and 59.4% for Monkey W. We further investigated whether the preference for the first information in delay1 affects the MD of the second information in delay2. This calculation was done separately for neurons with and without interaction effect (Fig. 4D). In the interaction group, the MD of second information was significantly higher if the first information is preferred (*p* = 3.13e-5 for pro-mode and *p* = 9.74e-7 for anti-mode). In contrast, neurons lacking an interaction effect did not exhibit a significant difference (*p* = 0.127 for pro-mode and *p* = 0.0614 for anti-mode). Overall, these results suggested that PMd maintains encoding of the firs information during the integration stage, yet there is a discernible interaction between reach and grasp.

**Figure 4.**
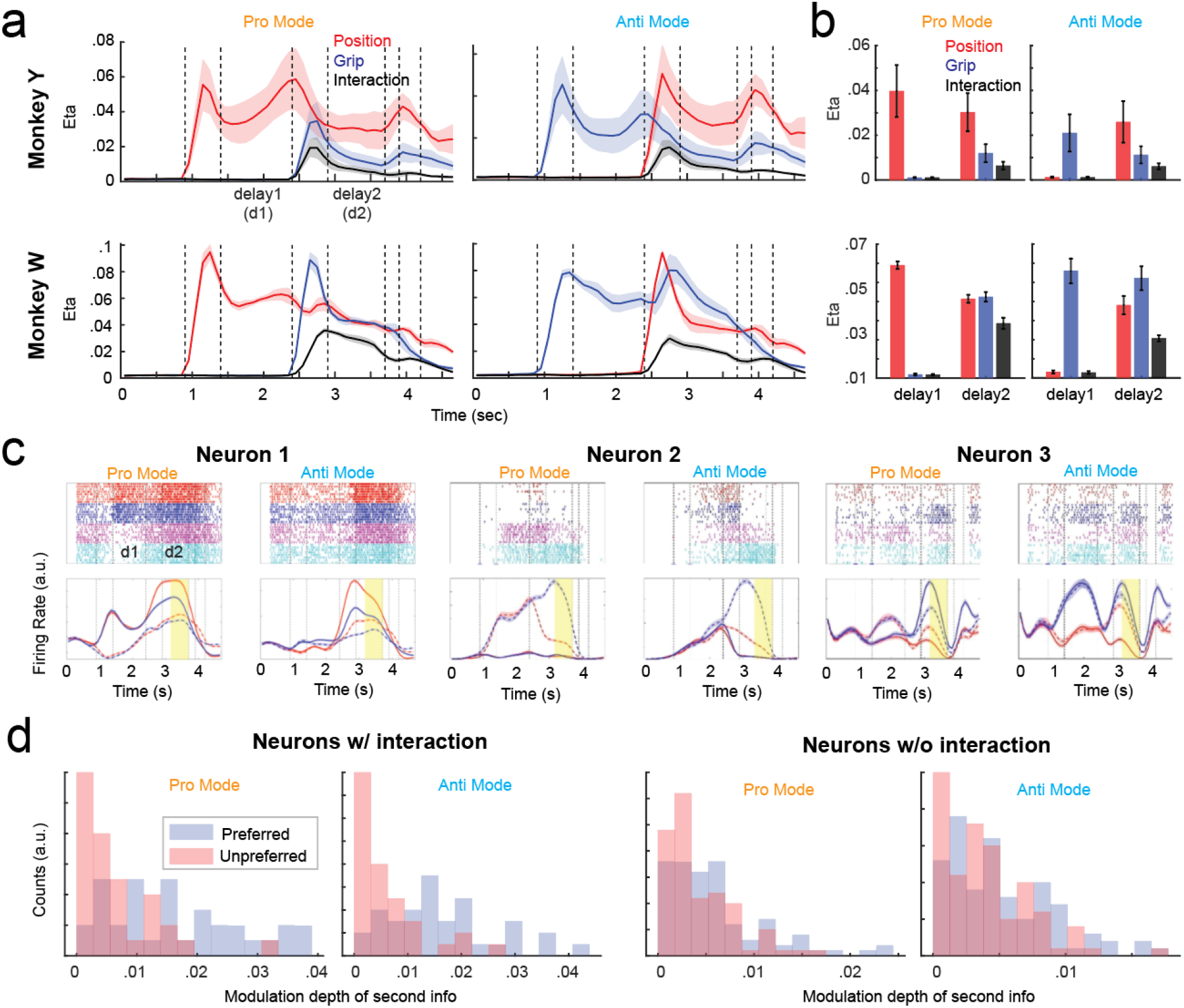
Reach and grasp are interactively encoded in PMd. **(a)** The eta values for position (red), grip (blue) and interaction (black) in ANOVA analysis are plotted against time during task. Data were averaged across all the sessions and shadow area shows standard deviation. **(b)** bar plot of eta value during delay1 and delay2 for both pro- and anti-mode. **(c)** Three example neurons show different interaction types (details in text). The conventions are the same as in Fig. 2a. **(d)** The histogram of modulation depth of second information during delay2 (i.e., grip in pro-mode and position in anti-mode) are plotted separately according to the preference of first information during delay1. The same analysis was applied for neurons with (left) and without (right) significant interaction effects.

### Population encoding structure for reach (grasp) was changed by grasp (reach)

To gain further insight into the way that reach and grasp interacted with each other in PMd, we scrutinized how encoding structure was changed by the second information at both population and single neuron level. To capture population encoding structure, we calculated coding direction (CD) for reach position in pro-mode and grip type in anti-mode. The CD is a multidimensional vector designed to maximumly differentiate the neural activity of two conditions (as detailed in the Methods section). We utilized CD as a quantitative metric of population encoding structure and calculated the cross correlation of CD across time in a trial to tract the changes (Fig. 5A). We revealed a low cross-correlation between the delay1 and delay2 epochs compared to the correlation within the delay1 or delay2 epochs themselves, which is highlighted by a square area in the heatmap. This low similarity between delay1 and delay2 was consistent across both monkeys and for both position coding in pro-mode and grip coding in anti-mode.

**Figure 5.**
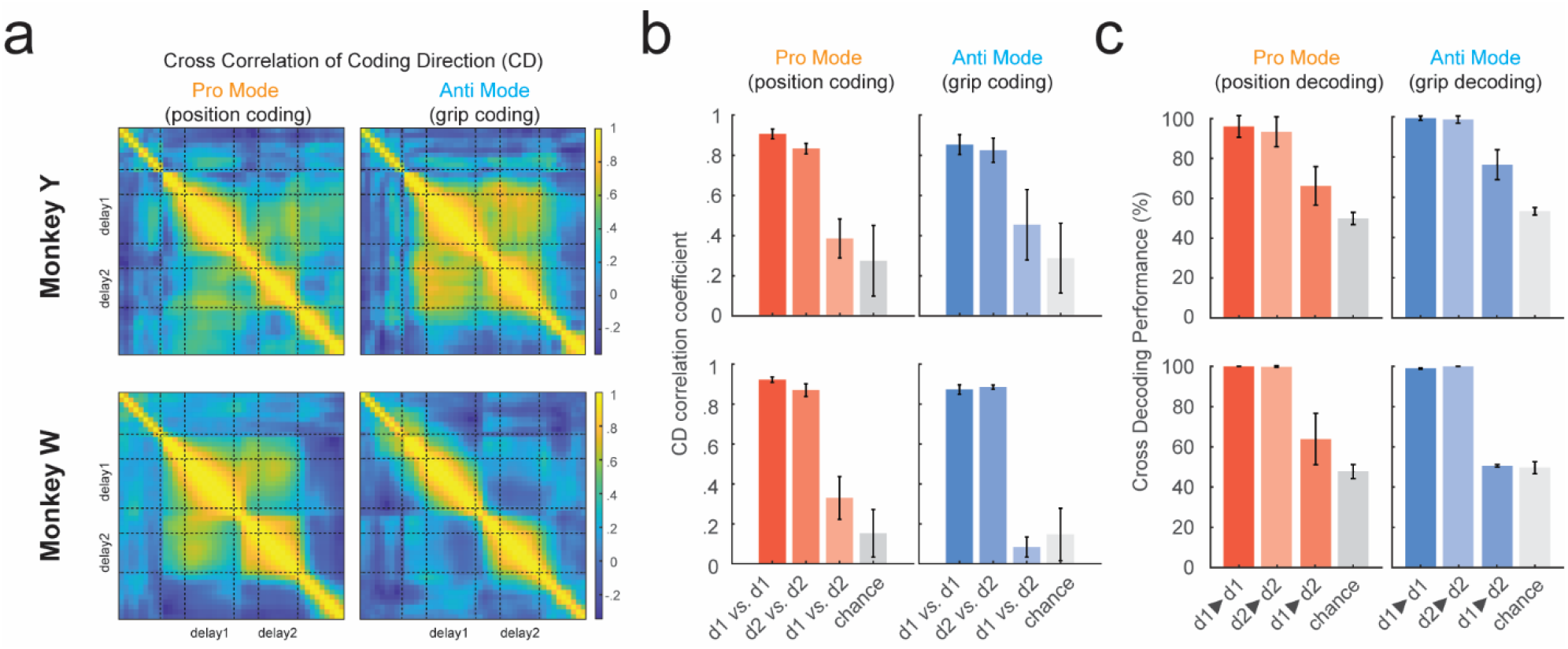
Population encoding structure for the first information changes when the second one arrives. **(a)** Cross correlation of coding direction (CD), a representation of population coding vector, for both position (pro-mode) and grip (anti-mode). **(b)** Averaged CD correlation coefficient for delay1 vs. delay1, delay2 vs. delay2, and delay1 vs. delay2. Chance level was calculated with shuffled data. **(c)** Same as (b) but for cross decoding performance. The items before and after arrow indicate training set and testing set, respectively.

The averaged correlation coefficient (cc) was quantified in Fig. 5B, indicating a significantly lower cc for delay1 vs. delay2 group. Cross single-trial decoding analysis further demonstrated that a decoder trained using delay1 activity achieved near chance-level performance for distinguishing trial types in delay2, confirming discernible patterns between delay1 and delay2 epochs.

At single-neuron level, we compared the MD between delay1 and delay2 epochs for each neuron. Neurons with significant modulation in at least one epoch were classified into four categories, as depicted in Fig. 6A: 1) stable, indicating no significant difference between delay1 and delay2; 2) reversed, where modulations in both delay1 and delay2 are significant but the preference is reversed; 3) enhanced, where MD in delay2 is significantly larger than in delay1; and 4) attenuated, where MD in delay2 is significantly lower than in delay1. The example neuron 1 (pro-mode), neuron 2 (anti-mode) in Fig. 4C and example neuron 2 (pro- and anti-mode) in Fig. 2A illustrated stable, enhanced and attenuated encoding, respectively. The example neuron in Fig. 6B illustrated the reversed encoding for grip type in both pro- and anti-mode. Across subjects and modes, only an average of 33.15% neurons manifested stable encoding, along with enhanced (20.33%), attenuated (30.03%) and reversed (16.49%) encoding patterns.

**Figure 6.**
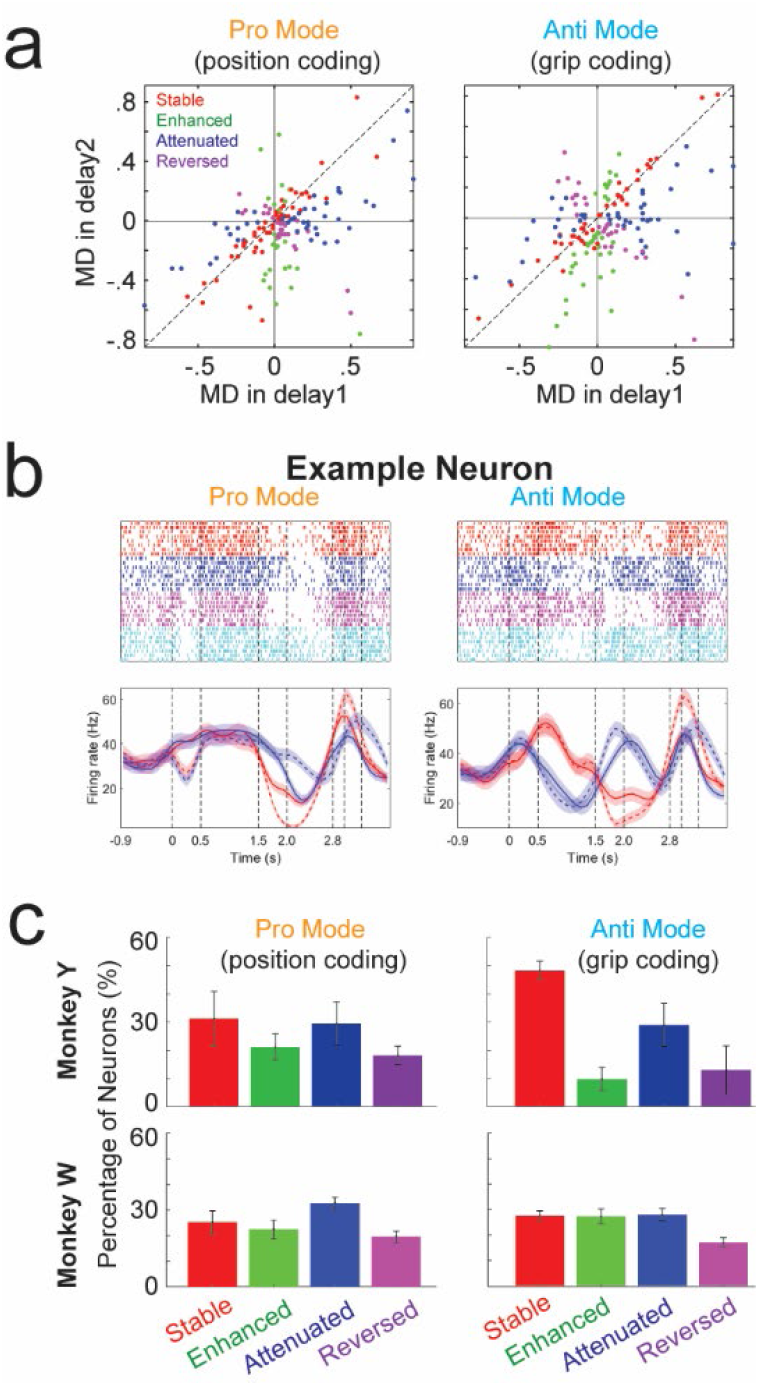
Encoding structure at single neuron level also changes. **(a)** Scatter plot of modulation depth in delay1 vs. delay2 for each significantly modulated neuron. Neurons are classified into four categories, including stable, enhanced, attenuated and reversed. **(b)** Example neurons showing the category of reversed MD. The conventions are the same as in Fig. 2a. **(c)** Percentage of neurons for each category in both pro- and anti-mode for both monkeys.

To dispel concerns regarding potential differences in cognitive states between delay1 and delay2, we also compared the two delay2 epochs of pro- and anti-mode, where the order of prompting is the sole difference (e.g., left power vs. power left). In this scenario, although an average of 48.57% neurons were stable, there were still a portion of neurons were enhanced (19.22%), attenuated (19.54%) and reversed (12.75%) as shown in Fig. S3. However, these differences began to dissipate at the later stage of delay2 epoch, and merged according to trial types just prior go cue (as depicted in Fig. S4). These results underscore that the encoding structure for the initial information underwent substantial changes with the addition of the second piece of information during the delay2 epoch, as evidenced at both the single neuron and population levels.

### CCA captures subspace for consistent encoding

The encoding structure for the first information prompted in SPRG task underwent significant changes with the introduction of the second information, yet the first information was persistently represented in the PMd, ensuring the monkey’s correct response. From neural dynamic perspective, the population representations during two delays take place in distinct yet overlapping neural subspaces. We hypothesize the existence of a low-dimensional latent space where consistent coding could be preserved over time. To investigate this, we utilized canonical correlation analysis (CCA) to identify the predominant co-variation patterns within PMd neural population. CCA identifies linear combinations or canonical variable pairs, of neural activities from delay1 and delay2 epochs that maximize the correlation between them.

Fig. 7A illustrated the session-averaged correlation coefficients (cc) for all CCA pairs. The highest cc reached up to 0.8, contrast to only 0.3 observed in shuffled data. An average of 75.0% and 65.4% (for monkey Y and W, respectively) of the pairs were significant (*Chi* test, p < 0.05) in each session; around the first eight pairs were particular higher than the rest, which could constitute of a space for consistent encoding. Fig. 7B displayed the activity of 4 example CCA components during dealy1 and delay2 epochs in both pro- and anti-mode. Taking CCA1 as an example, the activity distinguished between reach positions in pro-mode during delay1, and a similar pattern of separation recurred in delay2, indicative of consistent coding. This held true for other CCAs with high cc (like, CCA2 and CCA3), although the separation aperture could be less pronounced, as seen with CCA3. When the cc was low, such as with CCA30, the activity was more erratic, and the separation was not clearly discernible.

**Figure 7.**
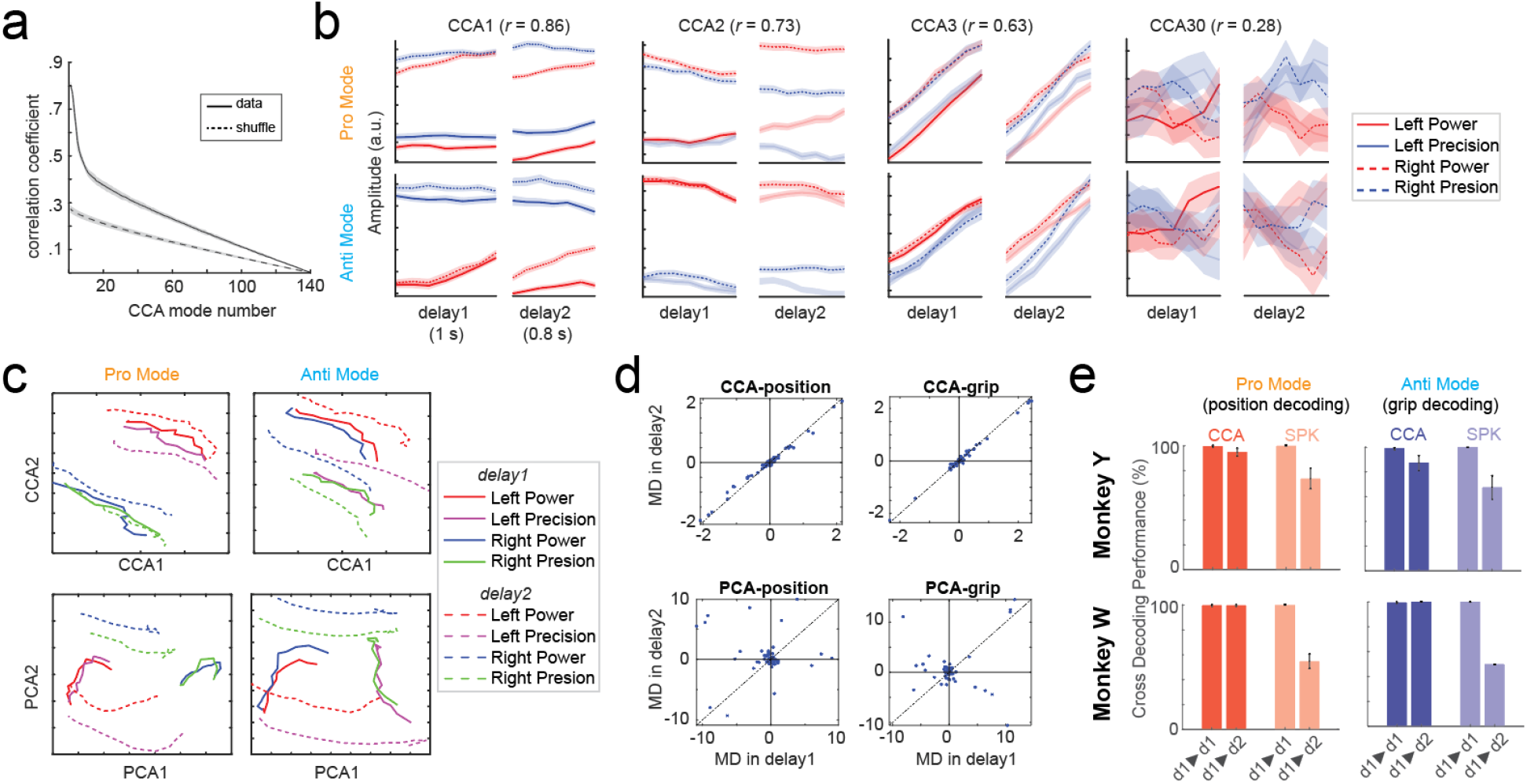
CCA subspace preserves consistent encoding. **(a)** Averaged correlation coefficient for each mode decomposed in canonical correlation analysis (CCA). Dashed line represents the same analysis using shuffled data. **(b)** The activities of four example CCA modes (CCA1, CCA2, CCA3 and CCA30) are plotted for delay1 and delay2 epochs. **(c)** The first two dimension of CCA (upper) and PCA (lower) are plotted for delay1 (solid lines) and delay2 (dash lines). **(d)** Scatter plot of neuronal modulation depth for delay1 vs. delay2 in CCA (upper) and PCA space. **(e)** Averaged decoding performance using both CCA activity and original spike activity (SPK). The items before and after arrow indicate training set and testing set, respectively.

To further visualize the CCA activities, we extracted the first the two CCA components to form a two-dimensional space and compared it with the PCA counterparts (Fig. 7C). Clearly, delay1 (solid) and delay2 (dash) activities could be classified with a single line for CCA. However, for PCA, the distribution of delay1 and delay2 were almost orthogonal to each other, necessitating two distinct lines for separation. We also quantified the MD for each CCA and PCA components between delay1 and delay2 epoch (Fig. 7D). The MD for CCA perfectly aligned with the right diagonal line for delay1 vs. delay2, in contrast with random scatters observed in PCA. Finally, the cross-decoding analysis revealed that CCA had significant above-chance decoding performance when using delay1 as training set and delay2 as testing set. These results confirmed that the encoding structures in CCA subspace for delay1 and delay2 epochs were consistent.

### Heterogeneous contribution to CCA

Next, we wondered how individual neurons contributed to the formation of CCA subspace. Given the stable (i.e., congruent) coding neurons reflect shared computation between delay1 and delay2 epochs akin to what CCA captures, it is plausible that these congruent neurons primarily contribute to the shared subspace.

To ascertain whether the shared subspace is primarily composed of congruent or incongruent neurons, we analyzed each neuron’s contribution to CCA. First, we quantified the congruence (i.e., stability) for each neuron by calculating the correlation between delay1 and delay2 epochs. Then we quantified each neuron’s contribution to the CCA (see methods). Fig. 8A presented the scatter plot of stability vs. contribution for each neuron, revealing no discernible relationship between these two metrics. We then sorted the neurons by ascending stability and plotted the contribution as a function of stability (Fig. 8B). The contribution values fluctuated without a clear trend as stability increased. As shown in Fig. 8C, neurons with both low and high stability exhibited similar distributions of subspace contribution.

**Figure 8.**
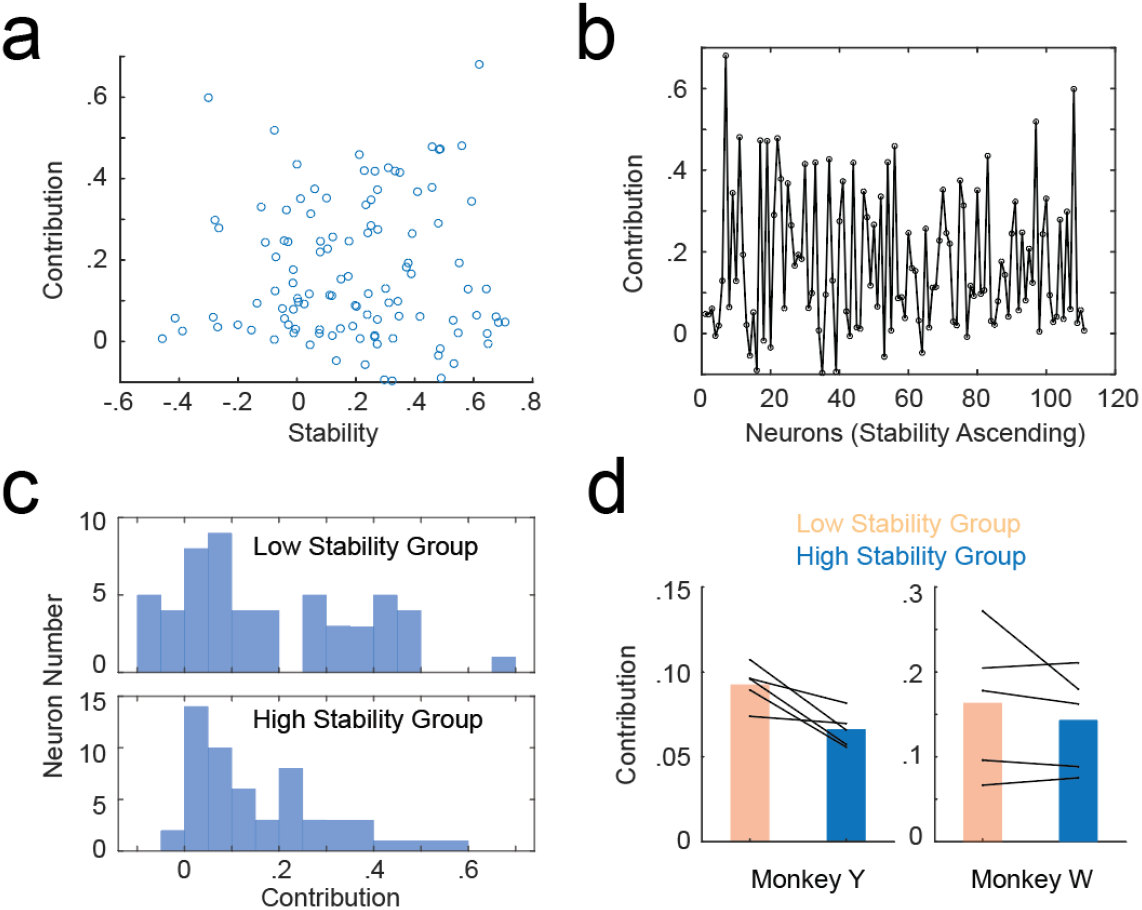
Neuronal contribution to CCA. **(a)** Scatter plot of contribution vs. encoding stability for each neuron in one example session. **(b)** The plot of contribution as a function of neurons with ascending stability. **(c)** Histograms of contribution for low (upper panel) and high (lower panel) stability group of neurons. **(d)** Session average of contribution for low and high group of neurons for Monkey Y (*p* = 0.3401) and W (*p* = 0.0222).

Quantitative analysis across sessions indicated similar (for Monkey W) or even lower contributions (for Monkey Y) for the high stability group (Fig. 8D). Essentially, both congruent and incongruent neurons contributed to the shared subspace.

To confirm that incongruent neurons do contribute to CCA, we simulated a group of neurons that exclusively composed of opposite modulation between delay1 and delay2 epoch (i.e., reversed neurons in Fig. 6). The MD of delay1 vs. delay2 was plotted in Fig. S5A, with example activity shown in Fig. S5B. We performed the same CCA analysis on this dataset and discovered that CCA could still capture a shared subspace, wherein components CCA1 and CCA2 displayed consistent MD (Fig. S5C) and activity (Fig. S5D). In summary, these results suggest that shared neural population structure is achieved by a heterogeneous ensemble of population, rather than being confined to a subpopulation of congruent neurons.

## Discussion

In this study, we investigated how PMd neurons sequentially integrate reaching and grasping information and plan for a successful reach-to-grasp movement. A novel task was designed to prompt serial cues for reach position and grip type with delays to guide the monkey through the grasping process. A significant proportion of neurons within PMd exhibited selectivity for both target position and grip type during the first delay epoch when only one type of information was presented. Upon the arrival of the second information, the PMd was observed to concurrently encode both types of information, a requirement for the successful execution of the task. However, the neuronal tuning property began to diverge from those observed in the first delay period. On one hand, some PMd neurons interactively encoded target position and grip type. On the other hand, a majority of neurons underwent a shift in their tuning properties following the introduction of new information, including enhancing, attenuating or even reversing the modulation rules established during the delay1 epoch. Despite these changes in tuning properties, congruent components could be extracted using CCA to form a stable subspace with consistent tuning. Moreover, the contribution of neurons to this stable subspace was not correlated with their individual tuning stability; incongruent neurons could significantly influence the stable subspace. These findings point to a sophisticated interplay between different types of information processing within PMd neurons during the motor planning phase.

### Cooperative coding in PMd for reaching and grasping movement

Over the past decades, the scientific community has amassed a more profound comprehension of the modulation of PMd neurons during reach-to-grasp movements. Early research was inclined to propose that the PMd was predominantly tasked with encoding reaching-related movements (Rizzolatti and Matelli, 2003; Tanné-Gariépy et al., 2002). Subsequent studies have uncovered that the PMd also partakes in encoding grasping movements (Stark et al., 2007; Takahashi et al., 2017), a revelation that points to the multifunctional modulation characteristics of the PMd. It is crucial, however, to acknowledge that neuronal coding patterns are intricately tied to the recording sites of electrodes in neurophysiological studies. According to Stark’s electrophysiological study (Stark et al., 2007), neurons implicated in reaching encoding are commonly situated near the superior precentral sulcus (SPcS), whereas those involved in finger movements are located at or below the arcuate sulcus (AS). Meanwhile, the PMd and PMv, as two subdivisions of the premotor cortex, are with their boundary delineated between the precentral hand and face motor areas, aligning with the frontal eye field (FEF) level on the dorsoventral axis (Stark et al., 2007).

In our experiment, the electrode arrays were strategically positioned above the AS bend (Fig. 1D), a region where reaching-related neurons were extensively recorded. For Monkey W, the placement of the electrode arrays was determined solely by empirical knowledge; whereas for Monkey Y, the electrode implantation was navigated by microstimulation to identify the mapping area of proximal limb movement. The outcomes from both monkeys exhibited analogous coding patterns concerning grasping movements (Fig. 1), potentially suggesting that neurons associated with grasping movements are generally dispersed above the AS bend. Moreover, we discovered that individual neurons within the PMd could concurrently encode target position and grip type during a period of multiple information representation, signifying their involvement in the motor planning of both reaching and grasping movements. This finding aligns with previous studies positing that reaching and grasping may overlap in both spatial and temporal dimensions (Begliomini et al., 2014; Verhagen et al., 2013) and further corroborates the perspective that reaching and grasping movements are not encoded separately in the primate brain (Takahashi et al., 2017).

A recent fMRI study has unveiled that the human motor cortex does not harbor a homunculus body map; instead, several body-part-specific zones are demarcated by integrative zones, blurring the lines of traditional discrete functional sectors (Gordon et al., 2023). This discovery was further substantiated by electrophysiology recordings (Jensen et al., 2023) and electrostimulation study in human (Roux et al., 2020), and is congruent with an earlier monkey study, where electrical stimulation at a single site of the motor cortex yielded coordinated multi-joint functional movements (Graziano et al., 2002). Our study, which demonstrates that reach and grasp are orchestrated within a singular brain region, harmonizes with these aforementioned investigations.

### Interaction between reaching and grasping encoding

Neuronal multifunctional information encoding, where a neuron responds to two or more types of information, typically involves information interaction (Follmann et al., 2018). That is, the modulation of one type of information significantly influences another. This interaction manifested in two primary aspects within our study.

Firstly, when a neuron simultaneously encodes target position and grip type, the impact of these two information types on the neuronal firing rate does not adhere to the principle of simple addition. For instance, a neuron may exhibit significant selectivity for grip type under conditions of a preferred target position, while maintaining only a basic firing rate for an unpreferred target position, irrespective of grip type. This could be attributed to the modulation of non-preferred information suppressing the modulation of another in some neurons. Neurons, which prefer only one or two specific position-gesture combinations, were extensively recorded in the PMd of both monkeys. This effect was quantified using *eta* values in ANOVA analysis, revealing a substantial portion of neurons engaged in interactive coding. We hypothesize that these neurons may be associated with the intricate cooperative encoding of reaching and grasping movements. As the monkeys’ proficiency in the behavioral task increases, the emergence of neurons tuned to specific position-gesture conditions may facilitate task completion with reduced effort.

On the other hand, neuronal information interaction is also evident in the modulation differences between periods of single-information encoding (delay1) and multi-information encoding (delay2). Upon the presentation of new stimuli, the majority of neurons in the PMd demonstrated significant changes in encoding the initial type of information, including substantial enhancement, attenuation, or even reversal of modulation depth. At the population level, the encoding of the first information was propelled into a new subspace, distinct yet not entirely orthogonal to the preceding one. This suggests that PMd neurons may employ two distinct encoding patterns during these two periods. Such temporal interactions have also been observed during sensory discrimination tasks (Rossi-Pool et al., 2017) and a decision making task (Banerjee et al., 2020) in previous studies, potentially indicating a universal principle for the temporal integration of information in goal-directed movements.

### Stable encoding preserved in a neuronal subspace

Although a substantial number of PMd neurons altered their encoding patterns following the introduction of new stimuli, both monkeys correctly performed actions during the experimental execution phase, indicating that the modulation of the initial information was retained in the PMd. One hypothesis remained that shared representations between the two delays occur at the level of neural population abstraction. In line with this hypothesis, the encoding patterns of the initial information remained stable before and after the arrival of new information within a specific subspace. Employing CCA, we projected the firing activities of all recorded PMd neurons from the two delay periods into a novel space (Gallego et al., 2020), where the encoding patterns displayed similar covariation among the neural population in a shared subspace that captures significant variance. This mathematical resemblance may underpin the stable encoding of movement in the PMd throughout the entire trial.

Previous neurophysiological studies either directly considered the modulation stability of individual neurons as their contribution to the stable encoding pattern of the neural network or selected neurons that positively impacted accuracy through neural decoding (Nuyujukian et al., 2014; Pandarinath et al., 2018). Generally, neurons with stable modulation characteristics were thought to contribute more to decoding accuracy than those with unstable characteristics. Contrary to previous studies, the contribution of individual neurons to the stable space was not significantly correlated with their own modulation stability; even neurons with reversal modulation characteristics could play a vital role in the stable space (Jiang et al., 2020). Mathematically, each common component in CCA is a linear combination of all input variables. The algorithm can transform unstable variables into stable ones by finding appropriate weights, thereby enabling unstable variables to positively contribute to the new space. From the perspective of brain neural networks, the activation pattern of individual neurons adjusts its weight and connectivity with other neurons. Thus, a neuron that prefers the left position during the single-information period might alter its connections within the entire neural network and exhibit a preference for the right position during the multi-information period.

### Limitations

Our experiments had two primary limitations. Firstly, the two delay periods, delay1 and delay2, as designed in our task, may serve distinct cognitive functions. For instance, the delay1 period might correspond to a working memory stage, while the delay2 period could be associated with sensory integration, working memory, and motor planning. The functional differences might result in divergent modulation patterns among individual neurons. To address this, we incorporated an analysis that exclusively compared the delay2 periods in both pro- and anti-modes, differing only in the prompt sequence, and the findings remained valid (Fig. S3). Secondly, Monkey Y was trained with a relatively rigid fixation to an LED, whereas Monkey W was equipped with an eye tracker to restrict ocular movement prior to the initiation of movement in the experiment. Consequently, it is challenging to differentiate whether the neurons in Monkey Y were encoding eye movements or motor planning, which could account for the higher prevalence of position-tuning neurons compared to gesture-tuning neurons.

In conclusion, this study offers further insights into how the encoding patterns of reaching and grasping may interact when temporally integrating two types of information to accomplish a final goal movement. Additionally, we have unveiled how the brain sustains a stable representation of information by identifying a stable subspace that is constructed by both congruent and incongruent neuronal populations.

## Methods

### Subjects

We trained two male monkeys (Macaca mulatta, Monkey Y: 7.4 kg, 4-year-old; Monkey W: 9.1 kg, 5-year-old) to perform a reach-to-grasp task (see below). After the monkeys mastered the task, we implanted two 96-channel microelectrode arrays (Blackrock microsystem, USA) into the arm areas of the primary motor cortex (M1) and the dorsal premotor cortex (PMd) in the hemisphere contralateral to their working hand. For monkey Y, ventral premotor cortex (PMv) was also implanted, but the data were not used in this study. We determined the implantation position of the microelectrode arrays based on the motor homunculus. For Monkey W, we used intracortical microstimulation to verify that the recording arrays were located in the forelimb region of the motor and premotor cortex. All surgical and behavioral procedures followed the Guide for the Care and Use of Laboratory Animals (China Ministry of Health) and received approval from the Animal Care Committee at Zhejiang University, China.

### Behavioral task

The behavioral task required the subject to sit in a customized chair with right hand fixed and perform a reach-to-grasp movement with left hand. An operation panel was placed in front of the subject, 20-25 cm away and at eye level, containing two identical handles (objects to be grasped), two white LEDs for position cue (left one for left object, right one for right object), an RGB LED for grip type (blue for power grip and yellow for precision grip) or go (green) cue and another white LED for fixation (see Figure 1A). The two handles were vertically fixed on the panel and separated by 15 cm. Each handle had four capacitive touch sensors installed inside to detect the bio-touch of four directions, respectively. The precision grip was indicated if only the front and back sensors were activated, while the power grip was indicated by the simultaneous activation of all four sensors. To minimize eye movement during the cue presentation phase, all LEDs were placed within a small area. Specifically, the fixation LED and the grip type /go LED were aligned on the vertical center line of the panel, 2 cm and 1.5 cm above the grasping object, respectively. The position LEDs were horizontally 1 cm apart from the grip type/go LED at the same height.

The behavioral task, sequentially prompted reach and grasp (SPGR) task, required the subject to respond to two action types sequentially, i.e., object positions (left vs. right) and grip types (power vs. precision). Therefore, the task had two modes depending on the order of the cues. In ‘pro’ mode, the position LED turned on first, followed by the grip type /go LED, with a delay in between. In ‘anti’ mode, the order was reversed. A trial began when the subject pressed a button down. At the same time, the fixation LED turned on to instruct the subject to fixate on it (baseline phase). The first action type cue appeared 900 ms after the trial started (cue1 phase: position cue in pro-mode; grip type cue in anti-mode). The first indicator LED lasted for 500 ms and then turned off. The trial entered the first delay phase (delay1), during which only the fixation LED was on. The second action type cue appeared 1000 ms after the first indicator LED turned off (cue2 phase: grip type cue in pro-mode; position cue in anti-mode). After another 500 ms, the second indicator LED turned off. The trial entered into the second delay phase (delay2), which lasted for 800-1200 ms. Then, the fixation LED turned off and the grip type/go LED turned green (go cue), signaling the subject to release the button and reach their hand to grasp the correct object with the cued grip type. The reaction time refers to the time interval from onset of go cue to the moment when the subject released the button. The trial was regarded as a failure if any of the following conditions occurred: 1. The subject released the button before the go cue. 2. The subject touched the wrong handle. 3. The subject grasped the handle with wrong grip types. 4. The subject’s gaze moved out of the predefined field of view (only of monkey W). In each session, the pro- and anti-mode trials were interleaved randomly at a trial-basis level.

### Neural recording and Data preprocessing

We used a commercial data acquisition system (Cerebus, Blackrock Microsystems, USA) to record and preprocess the continuous neural activities. The analog signal from the microelectrode array was amplified, bandpass filtered (Butterworth, 0.3–7500 Hz), digitized (16-bit resolution, 30 kHz sample rate) and highpass filtered (Butterworth, at 250 Hz) sequentially. Spike activities were detected from the filtered signal by applying a threshold of -5.5 times the root mean square (RMS) of the baseline signal, and then sorted using Offline Sorter (Plexon Inc., USA). The sorted spikes were binned with a nonoverlapping time window of 10 ms and smoothed with a Gaussian kernel with stand deviation of 50 ms to construct peristimulus time histograms (PSTH). We calculated the 95% confidence interval across trials in each condition to show the variance. The variance of neural population in PMd is calculated from the Gaussian-smoothed neural activity across all trials and all neurons.

### ANOVA

We quantified the neuronal response to specific stimulus using the ANOVA method. Considering two types of information (reach position and grip type) were involved in the experiment, both one-way and two-way ANOVA were applied to investigate neuronal modulation to single information and multiple information, respectively. We first got firing rates in the first 800 ms of the delay period and calculated the p value of each individual data segment in Matlab.

We designed a quantitative index η(*eta*) based on the two-way ANOVA to compare the modulation depth of a neuron to different stimuli. The sum of squares (SS) in ANOVA reflects the effect of a factor on the variable, and here we normalized it with the total SS to quantify the response level of a neuron to a stimulus.

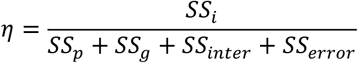

where *SS*_*p*_, *SS*_*g*_, *SS*_*inter*_ and *SS*_*error*_are the sum of square of target position, grip type, interaction and error in two-way ANOVA. *SS*_*i*_ are selected from *SS*_*p*_, *SS*_*g*_, and *SS*_*error*_. In Fig 4a, *eta* is calculated used a sliding window of 50 ms along the trial. Eta indicates the response degree of a neuron to a specific stimulus or interaction, and varies between 0 (the neuron does not tune to the stimulus at all) and 1 (the neuron strictly and exclusively responds to the stimulus) theoretically. Due to the uncertainty of spikes and the recording error of neural signals, *eta* cannot be equal to 0 or 1 exactly.

### Modulation depth

Modulation depth (MD) measures the neuronal firing activity in response to specific stimulus or action. We calculated MD in a predetermined time window for reaching and grasping movement individually. MD was defined as the normalized difference in average firing rates between two action choices of the same movement type.

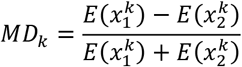

where k indicates the movement type (reach position or grip type), 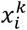 indicates the firing rate of specific action choose for k action type (left or right object for reach position; power or precision grip for grip type).

*E*(·) means calculating the average firing rate across trials in the predetermined time window.

### Decoding Analysis

We used support vector machine (SVM) to decode underlying motor intention from activities of all recorded neurons to investigate the modulation of neural cluster in PMd. The Gaussian-smoothed neural activity was first binned with a time window and the firing rate was averaged in this window for individual trials and neurons. Then all trials were divided into two or four groups (two groups for single information classification, four groups for dual information classification. We performed a binary or four class classification to investigate whether the neural activity encode the target information. All trials are randomly grouped into 10 groups to make a 10-fold cross validation test.

### Coding direction

We calculated coding direction (CD) to investigate the tuning property of neural population (Li et al., 2016) in PMd. The CD index is a vector that represents the differences in firing rates between two stimulus conditions for all neurons. The CD for reach position is defined as follows:

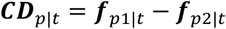

where *f*_*p*1|*t*_ and *f*_*p*2|*t*_ are vectors including the average firing rates of all neurons in a time window for left and right positions, respectively. Similarly, the coding direction of grip type was calculated as:

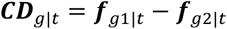

where *f*_*g*1|*t*_ and *f*_*g*2|*t*_ are the average firing rate vectors of all neurons for power and pinch grip, respectively.

To examine how the tuning property of PMd neural population changes over time, we calculated the coding directions (CDs) sequence using a 200-millisecond time window throughout the entire trial. Subsequently, we computed the self-correlation matrix of this CD sequence as depicted in Fig. 5. Each element in the matrix represents the correlation of populational tuning property between two specific time points.

### Neuronal encoding change types

Neurons with selectivity for target position or grip type widely changed their modulation after new stimuli appeared. Therefore, we divided all selective neurons into four groups (i.e., stable, enhanced, attenuated, reversed) by comparing their individual modulation for a type of information (target position or grip type) between delay1 and delay2 periods. Two 500-millisecond time windows, which started at 300ms after the cue light turned off, were used to extract neural activities in delay1 and delay2 period, respectively.

First, we use ANOVA to assess the significance of neuronal selectivity for specific information in two delay periods, respectively. All PMd neurons are divided into four groups including “silent”, “active”, “appeared” and “disappeared”. The “active” and “silent” neurons show significant selectivity and no selectivity in both delay periods, respectively. The neurons with significant selectivity in delay1 but no selectivity in delay2 were defined as “appeared” neuron, while the neurons with no selectivity in delay1 and significant selectivity in delay2 are regarded as “disappeared” neuron.

Second, we further divided “active” neurons into four groups including “stable”, “enhanced”, “attenuated” and “reversed”. Suppose 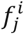 and 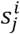 are average and variance of firing rates across trials for *ith* movement choice (for position: left, *i* = 1, right, *i* = 2; for grip type: power grip, *i* = 1, precision grip, *i* = 2) in *jth* delay period, where *i, j* ∈ [1,2]. 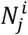 is the trial number for *ith* ovement choice in delay period. In our experiment, the trial number for individual movement choice are equal, so we omit the superscript of 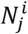 to simplify the notation. The average and variance of modulation depth in *jth* delay period, *m*_*j*_ and *s* _*j*_, are calculated as

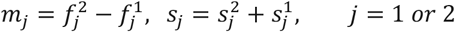

We employed a two-sample t-test to assess the significance of the difference in neuronal modulation between two delay periods. The t-value of t-test was calculated as

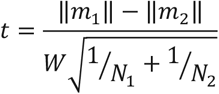

Where *W* are defined as

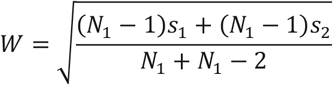

Then we use a t-distribution table to find the p-value *pp* that corresponds to the t-value and degrees of freedom. All “active” neurons are divided into four groups (“stable”, “enhanced”, “attenuated” and “reversed”) according to their t-test results and modulation depths in two delay periods.

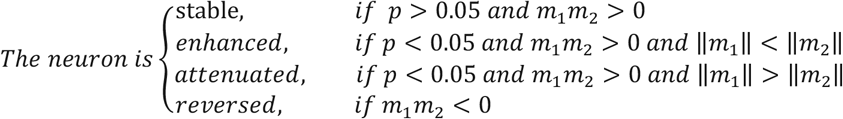

Finally, considering that the “appeared” and “disappeared” neurons are the extreme cases of “enhanced” and “attenuated” neurons, respectively. We added “appeared” and “disappeared” neurons into “enhanced” and “attenuated” groups, respectively.

### Stability index of tuning property

To examine the changes of neuronal coding patterns across trial phases, we measured the modulation stability of individual neurons in delay1 and delay2 phases using Pearson correlation coefficient.

Specifically, the binned spike trains for all action conditions are concatenated trail by trail and smoothed with 50ms Gaussian kernel in the two delay phases separately. The stability index of a specific neuron is defined as the Pearson correlation coefficient of between its firing activities in two phases. The stability indexes of all neurons were sorted in descending order to obtain a gradient table of modulation stability.

### Canonical correlation analysis

To investigate the correlation of neural activities between delay1 and delay2 phase, we use canonical correlation analysis (CCA) to extract the similar components of the two phases (Gallego et al., 2020).

Unlike principal components analysis based on the principle of maximizing the projection variance, CCA finds the projection direction of two variables to maximize the correlation between the projected data.

Given two T by M matrices *X*_1_ and *X*_2_ (T and M are numbers of samples and neurons, respectively) as the neural activities in delay1 and delay2 phases, CCA projects them into a new manifold with two linear transformation matrices so that the projected activities have maximal correlation.

CCA can be calculated using singular value decomposition of matrices. First, we obtain the transformation matrices *Q*_1_ and *Q*_2_ using QR decomposition, *X*_1_ = *Q*_1_ *R*_1_, *X*_2_ = *Q*_2_*R*_2_. The first M columns of *Q*_1_ and *Q*_2_ provide an orthonormal basis for each column of *X*_1_ and *X*_2_. Then a singular value decomposition is applied to the inner product matrix of *Q* _1_ and *Q* _2_.

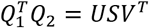

The elements of the diagonal matrix *S* are the canonical correlations, which are sorted from large to small. The projection matrices that project *X*_1_ and *X*_2_ into a new manifold can be calculated as follows.

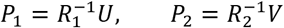

We concatenated the spikes trial by trial to build the neural activity metrices *X*_1_ and *X*_2_ in delay1 and delay2 phases, separately, and fed them into the CCA function. The projected neural activities *Y* _1_ and *Y* _2_ can be calculated with the projection matrices *P*_1_ and *P*_2_.

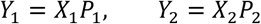

Each dimension of the projected activities could be regarded as a neuron, from with the PSTH, modulation depth and movement decoding were calculated (Fig 7).

### Contribution index for new CCA manifold

In CCA, each dimension in the new manifold is a linear combination of the firing rates of all neurons and neurons contribute variously to specific dimension. The contribution of a neuron to the new manifold is generally measured by correlation. Specifically, the contribution of ith neuron in delay1 phase is defined as follows (Bae et al., 2020):

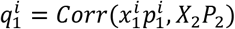

Where 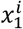 and 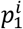 represent firing activity and projection weight of ith neuron in delay1 period, which are parts of *X*_1_ and *P*_1_, respectively. A high value of 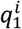 implies 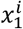 plays a significant role in the relation between *X*_1_ and *X*_2_. Similarly, we computed the contribution of ith neuron in delay2 period to the new manifold.

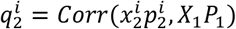

Then the integrated contribution index of ith neuron is defined as the average of 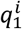 and 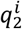.

To figure out whether neurons with stable modulation contribute more to the new manifold in the CCA, we investigated the correlation between stability and contribution index. In Fig 8b, the neurons were sorted in descending order in horizontal axis and their corresponding contribution index are shown in vertical axis. In Fig 8c, we divided neurons into two groups according to the stability index and compared their histograms of contribution indexes.

### CCA simulation

We employed an extreme simulation method to investigate whether neurons with unstable modulation also contribute in CCA. Specifically, we simulated the firing activities of 100 unstable neurons in delay1 and delay2 phases. Those neurons exhibited modulations that were either enhanced, attenuated or reversed. Our simulations encompassed all eight action choices involved in the behavioral task, with each choice consisting of 50 trials. The firing rates of these simulated neurons are linearly correlated with reaching position and grip type. Notably, there was no interaction between these two types of information. The firing rate of each simulated neuron was defined as follows:

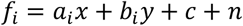

where x and y represent reaching position and grip type, respectively. Their values were chosen from two options: 1 (indicating left position or power grip) and 2 (indicating right position or precision grip). *a*_*i*_ and *b*_*i*_ represent weights in delay *i* phase with respect to reaching position and grip type, respectively. Their values, ranging from -10 to 10, represented the preferred choice and modulation depth of a simulated neuron. To simulate unstable neuron, the values of weights were intentionally set differently between delay1 and delay2 phase. For instance, if a neuron has a_1_ = 10 and a_2_ = 0, it means the neuron has an attenuated modulation with respect to reaching position.

### Laplacian eigenmaps

Laplacian eigenmaps (LapEig) is a graph-based dimensionality reduction method, which projects the high-dimensional data into a low-dimensional space while preserving the distance graph of the raw data (Belkin and Niyogi, 2003). Given a dataset of *n* samples, the LapEig method first constructs an adjacency map by treating each sample as a graph node and compute edges from the root node to its k nearest neighbors. The edge weight *w*_*i,j*_ between two nodes *x*_*i*_ and *x*_*j*_ is defined as

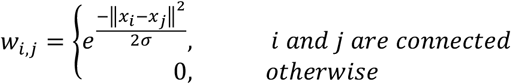

Where *σ* represents the scale factor. This function ensures that nodes close to each other have larger edge weights. The adjacency matrix of the dataset *W* is composed of all edge weights *w*_*i,j*_ between two nodes and the degree matrix *D* is a diagonal matrix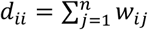. In order to project the original data node *x*_*i*_ into *d*-dimensional space node *y*_*i*_ while preserving neighboring relationships, the objective function of LapEig is defined as

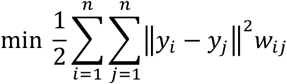

The optimization problem can be deduced to the following form:

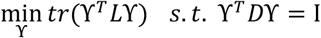

Where *L* is the Laplacian matrix that is defined as the difference between the degree matrix and the adjacency matrix. The ith row of *ϒ* is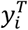. The constraint defines the scale of the solution and eliminates trivial solutions such as all zero points. The deduced problem can be solved by computing the first *d* eigenvectors of *L* because *L* is usually sparse.

## Acknowledgements

This work was supported by STI 2030—Major Projects (2022ZD0208900), National Natural Science Foundation of China (62336007), Pioneer R&D Program of Zhejiang (2024C03001), the Starry Night Science Fund of Zhejiang University Shanghai Institute for Advanced Study (SN-ZJU-SIAS-002), and the Fundamental Research Funds for the Central Universities (2023ZFJH01-01, 2024ZFJH01-01). We thank Ms. Guihua Wan for assistance with animal care, handling, and training, Dr. Junming Zhu for surgical procedures.

