## Supplemental Figures for "Interactively Integrating Reach and Grasp Information in Macaque Premotor Cortex"

### Supplement Information

**Authors**:

Junjun Chen[1,2], Guanghao Sun[1,4], Yiwei Zhang[1,4], Weidong Chen[1], Xiaoxiang Zheng[1,4], Shaomin Zhang[1,3]*, Yaoyao Hao[1,3,5]*

**Affiliations**:

1 Qiushi Academy for Advanced Studies, Zhejiang University, Hangzhou, 310027, China

2 School of Rehabilitation Sciences and Engineering, University of Health and Rehabilitation Sciences, Qingdao, 266114, China

3 The State Key Lab of Brain-Machine Intelligence, Zhejiang University, Hangzhou, 311100, China

4 Department of Biomedical Engineering, Zhejiang University, Hangzhou, 310027, China

5 Nanhu Brain-computer Interface Institute, Hangzhou, 311100, China

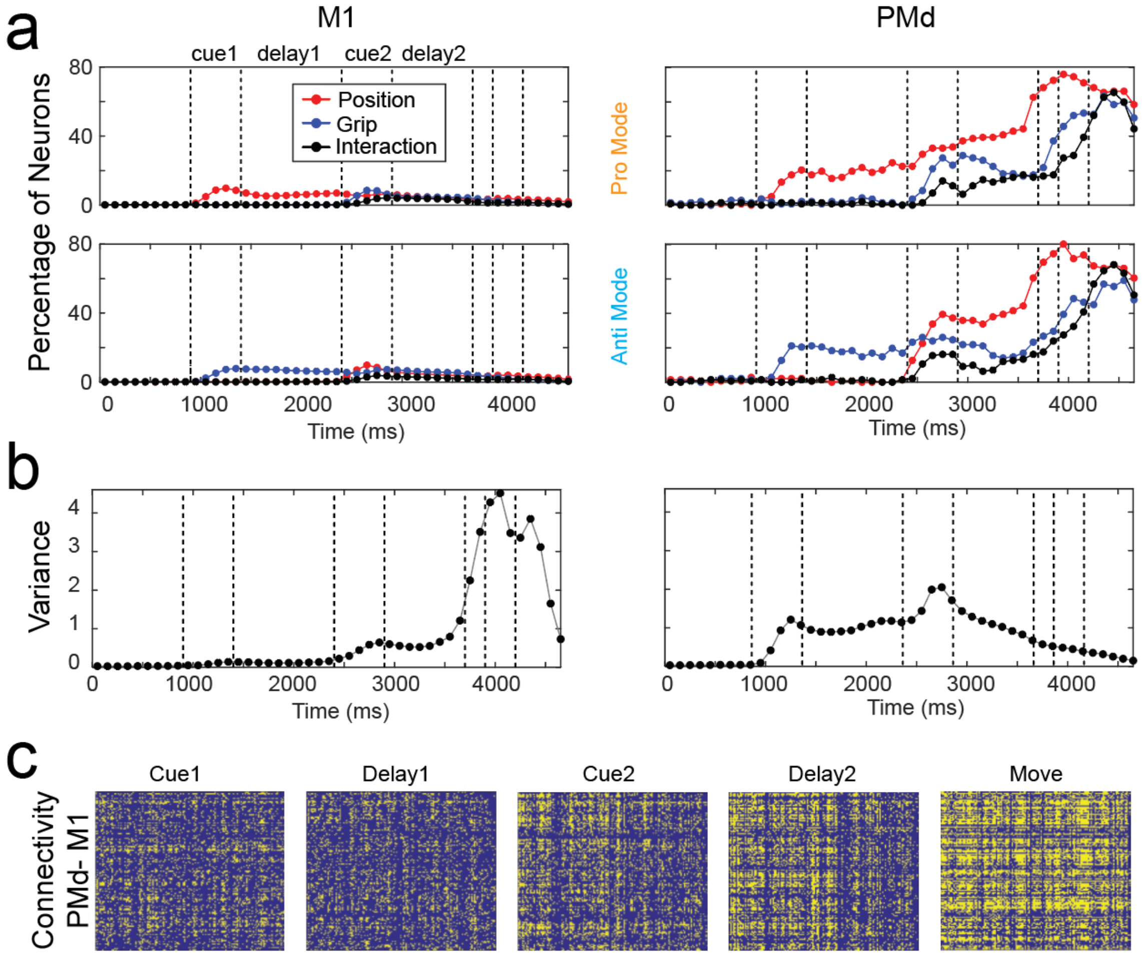


**Supplementary Figure 1. Comparison of neural properties in PMd vs. M1. (a)** Percentage of neurons that significantly modulates position (red), grip (blue) and interaction (black) in both pro (upper panel) and anti (lower panel) modes. **(b)** Instantaneous total population variance as a function of time. **(c)** Connectivity matrix between neurons in PMd and M1. Yellow color indicates significant connection while blue indicates no significance.

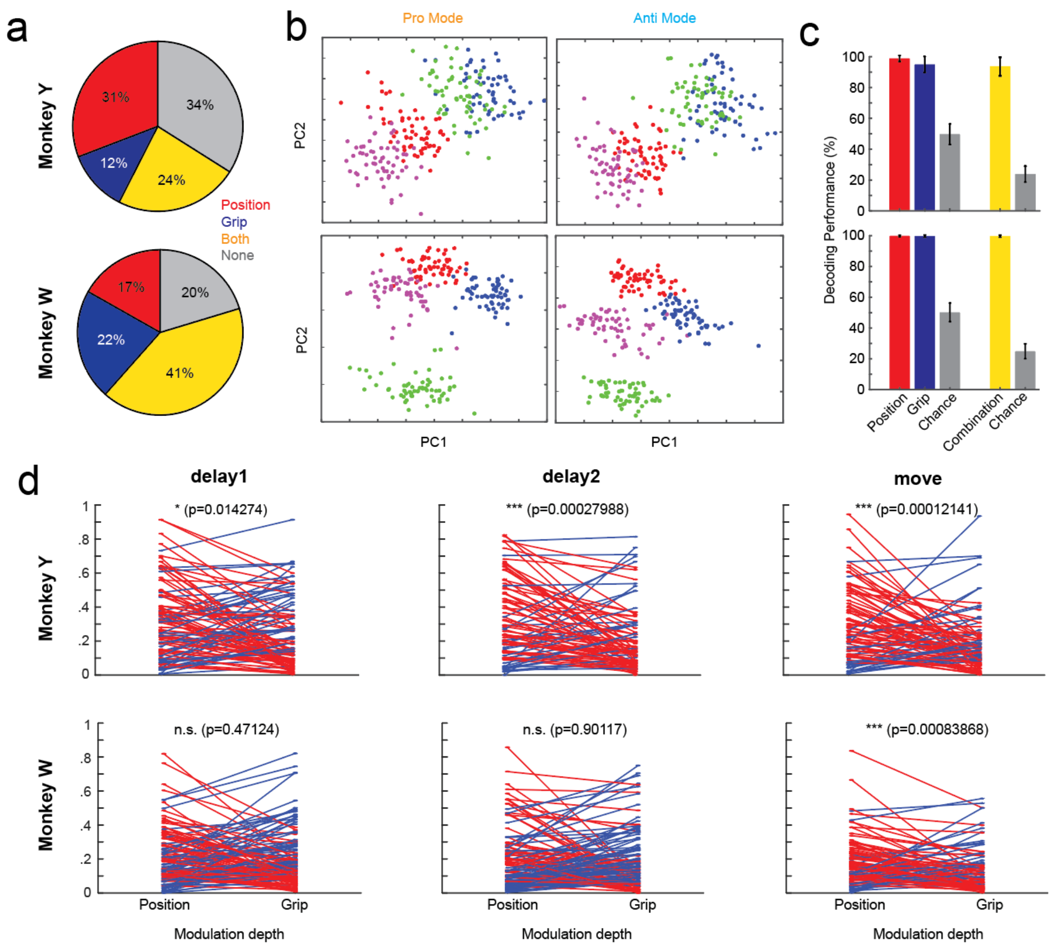


**Supplementary Figure 2. PMd encodes both reach and grasp in delay2. (a)** Percentage of neurons in PMd that modulate position (red), grip type (blue), both (orange) and neither (gray) during delay2. **(b)** Neural activity during delay2 was dimensional reduced to 2D space (PC1 vs. PC2) using principal component analysis (PCA). Each dot indicates averaged activity in one trial. **(c)** The decoding performances for position (red), grip (blue) and combination (yellow) using delay2 activity. The gray bar indicates chance level using shuffled data. **(d)** Comparison of modulation depth (MD) for position and grip for each neuron (each line). Red indicates higher position MD and blue indicates higher grip MD. Asterisk indicates significant level and n.s. means not significant.

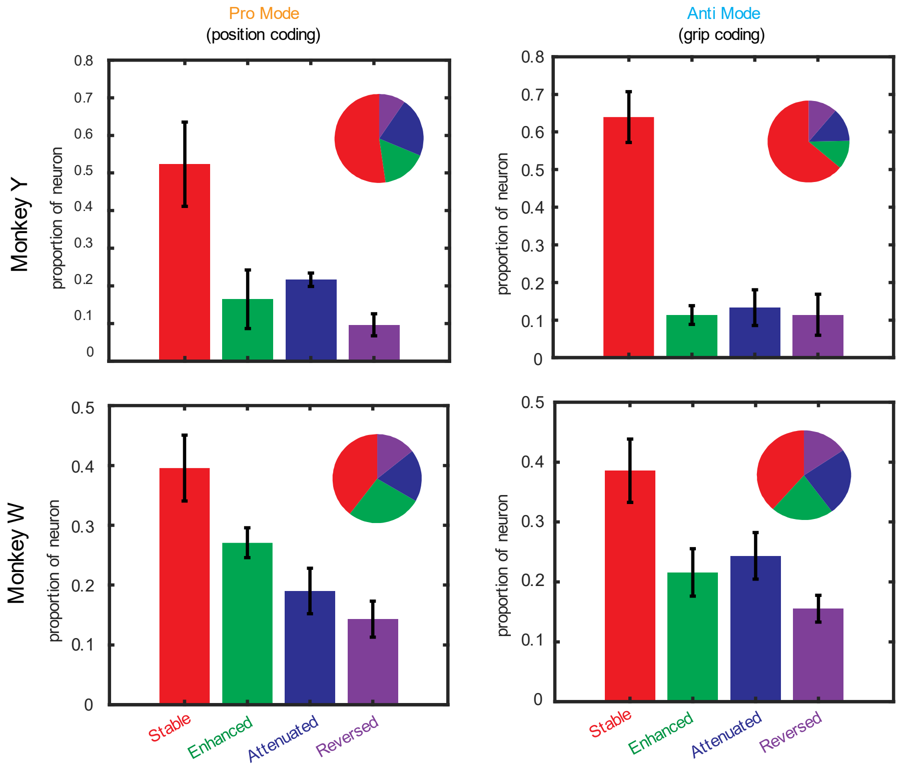


**Supplementary Figure 3. Encoding structure for Pro vs. Anti-mode in delay2.** Percentage of neurons in delay2 for each category (stable, enhanced, attenuated and reversed) in both pro- and anti-mode. The bar and pie charts represent the same data.

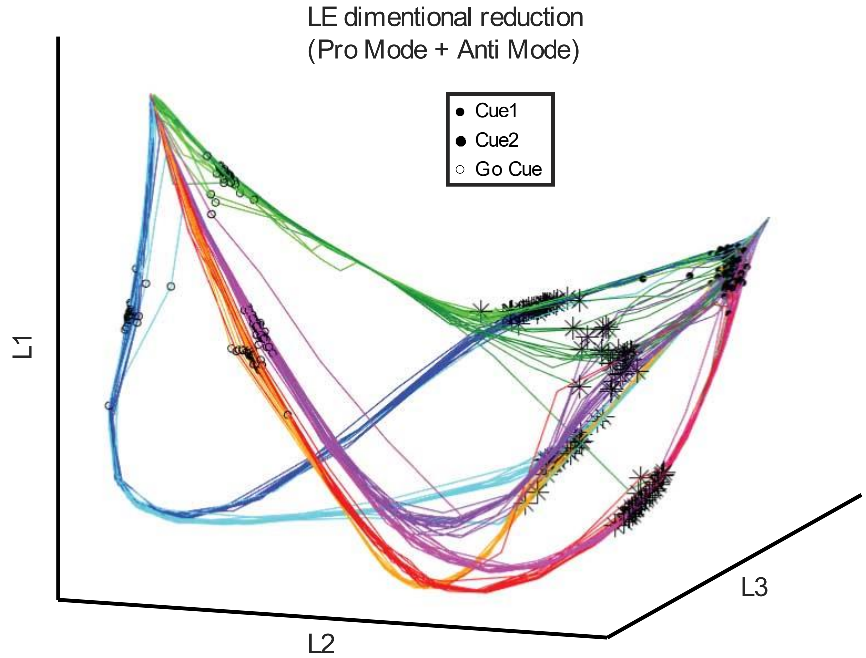


**Supplementary Figure 4. Neural activity after LE dimensional reduction.** The neural activity in one example sessions was reduced to 3-dimension using Laplacian eigenmap (LE). Different colors represent the four conditions (i.e., left power, left precision, right power, and right precision) in both pro- and anti-modes (8 in total). Dot, asterisk and circle indicate start of Cue1, Cue2 and Go Cue, respectively.

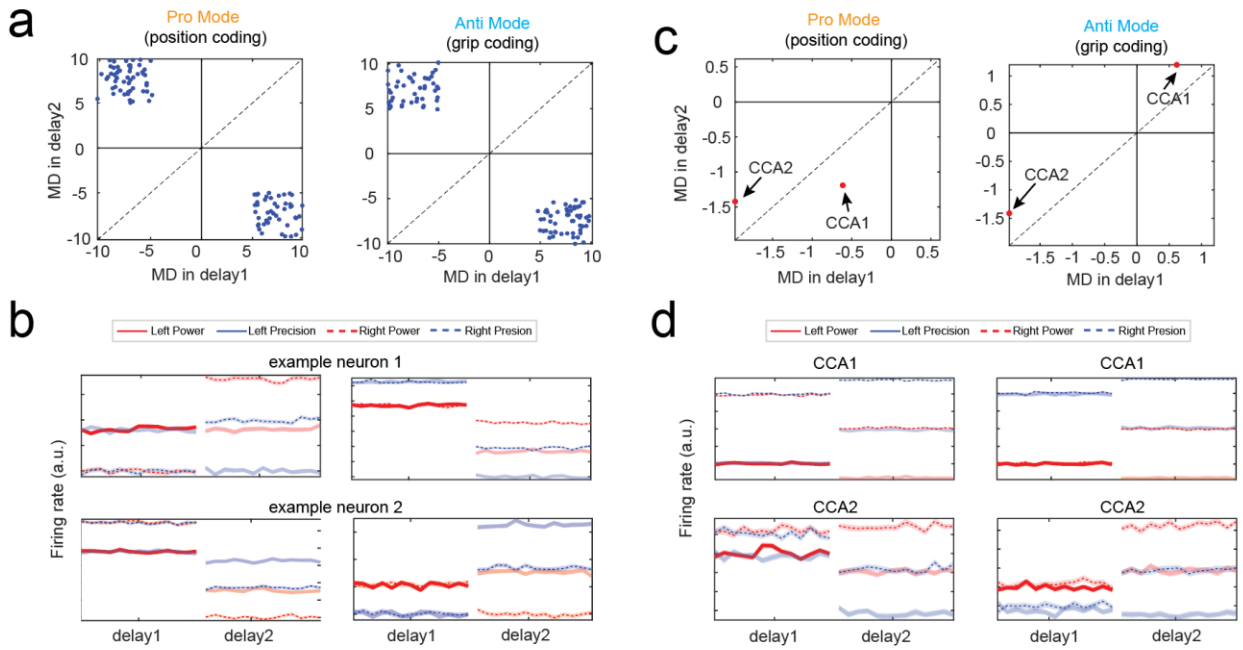


**Supplementary Figure 5. Simulation analysis of CCA. (a)** The modulation depth (MD) of a group simulated neurons which have opposite modulations in delay1 vs. delay2. Each dot represents one neuron **(b)** two example neurons from (a) showing activities in both pro (left) and anti (right) mode. **(c)** The MD in delay1 vs. delay2 for the first two CCA dimension. **(d)** Same as (b) but for activity of the first two CCA dimension (CCA1 and CCA2).
